# Structural thermodynamics by cryo-EM resolves minute energy differences in heterogeneous protein complexes

**DOI:** 10.64898/2026.10.07.756995

**Authors:** Lars Boyens-Thiele, Maša Bićanin, Kasper Engholm-Keller, Andreas Bøggild, Thomas Boesen, Birte Svensson, Alexander Kai Buell, Jens Preben Morth

## Abstract

Hetero-oligomeric protein complexes play crucial roles across biology, providing combinatorial diversity and context-specific functionality in membrane receptors, molecular chaperones and beyond. Yet, existing methods are limited in their ability to resolve and quantify the assembly energetics of complex biomolecular populations. Here, we introduce a comprehensive method combining iterative 3D classification, error-rate quantification via simulated particles, and global thermodynamic modeling to extract interaction energies from cryo-EM data. Using hexameric cruciferin (the primary rapeseed storage protein) as a model, we reconstruct a 1.83 Å structure and distinguish three highly similar 50 kDa isoforms. We resolve the complex compositional heterogeneity of the hexameric species at the single-particle level and fit a global binding model revealing pairwise interaction energies between isoforms differing by fractions of the thermal energy (*k_B_T*). This method offers a powerful, broadly applicable tool to characterize the compositional heterogeneity and energetics of biomolecular complexes.

## Introduction

Many proteins function as large, compositionally heterogeneous complexes, requiring structural and energetic characterization of their assembled states to fully understand their biological roles^1^. Diverse examples of bio-molecular complexes showing compositional heterogeneity include molecular chaperones and their cochaperones^2,3^, membrane-bound receptors^4^, and seed storage proteins^5,6^. In order to quantify the energetics of such heterogeneous assemblies, the populations of all possible assembly states need to be quantified. Ensemble methods, such as NMR spectroscopy^7^ and X-ray crystallography are often unable to resolve diverse complexes in populations with high combinatorial diversity. Most single molecule/particle methods are limited by the need for fluorescent labeling (e.g., smFRET^8^) or their intrinsic resolution (e.g., mass photometry^9^ and native mass spectrometry^10^). Single-particle analysis (SPA) cryo-electron microscopy (cryo-EM) uniquely allows access to both structural and population information on large biomolecular complexes^11,12^. By combining signals originating from many single particles, SPA offers unique possibilities for characterizing the structural heterogeneity of macromolecules^13^. The heterogeneity may either be conformational heterogeneity arising from chemically identical species adopting distinct conformations, or compositional heterogeneity resulting from the coexistence and assembly of chemically distinct species^14^. In recent years, several algorithms have been developed for investigating systems displaying continuous conformational heterogeneity^15–19^. For the characterization of populations of discretely distinct species, such as those arising from compositional heterogeneity, the method of 3D classification has been developed for unsupervised classification of particles based on their structural similarities^20^. Focused 3D classification has been applied to describe populations of diverse heterogeneous biomolecules^4,21–23^.

Cruciferin, the predominant seed storage protein (SSP) of rapeseed (*Brassica napus*) is an 11S globulin making up around 60% of the protein content of the seeds by mass^24^. Similar to other 11S globulin (legumin-like) SSPs, cruciferin belongs to the cupin superfamily, having a shared fold composed of two β-barrels per monomer^25^ and forms a 300 kDa hexameric assembly^26,27^. Multiple cruciferin isoforms with high sequence similarity occur in rapeseed and are thought to form heterogeneous hexameric complexes, although the exact nature and diversity of the heterohexamers remain to be characterized^27,28^. In homologous systems, a variety of of co-assembly specificities have been observed, ranging from the formation of highly specific co-assemblies^6,29^ to systems unable to assemble into crystals, the latter case necessitating the use of single-isoforms mutants to achieve crystallisation^30^.

Here we present a detailed structural and thermodynamic analysis of rapeseed-derived cruciferin. The cryo-EM structure of hexameric cruciferin is solved at 1.83 Å resolution. By applying symmetry-expansion, signal subtraction, and focused 3D classification, we identify and distinguish three isoform classes and determine the populations of all possible hexameric assembly states. By quantifying misclassification rates, we correct the cryo-EM-derived population estimate and achieve results in agreement with the overall composition as quantified by LC-MS. We further quantify the relative energetics of cruciferin hexamerization by the different isoforms and find that the energies of the different hexameric arrangements differ by fractions of the thermal energy, *k_B_T*. Cruciferin represents an extreme case of compositional complexity and promiscuity, and thus serves as a rigorous test case for population and thus energetic analysis by advanced single-particle cryo-EM methodologies. We define ‘structural thermodynamics’ as the quantification of interaction energies from a distribution of structural states observed via cryo-EM. This framework provides a generally applicable method to characterize the compositional heterogeneity of biomolecular complexes and will enable a comprehensive understanding of assembly diversity and energetics of native protein complexes across a wide range of biological systems.

## Results and discussion

### A high-resolution cryo-EM structure of hexameric cruciferin

We set out to analyse the energetics of cruciferin assembly through high resolution structural analysis of the assembly states. A cryo-EM density map of hexameric *B. napus* cruciferin was reconstructed at 1.83 Å, by imposing *D*_3_ symmetry (Schoenflies notation^31^), hence forcing homogeneity of the six constituent subunits (EMD-58942). Fourier Shell Correlation (FSC) and Guinier plots for the reconstruction are shown in Supplementary Fig. 1. Homohexameric structures of three cruciferin isoforms (UniProt gene names CRU1, BnC1, and CRU4) were refined against the same reconstructed density map. The reconstruction metrics for the three refined structures are given in Table 1. These three cruciferin isoforms were chosen as representative sequences spanning the diversity of previously annotated sequences in UniProt^32–37^. Cartoon representations of the refined homohexameric structure of CRU4 are shown in Fig. 1a, illustrating the six-fold (*D*_3_) symmetry of the assembly. The overall structure of the cruciferin complex closely matches the classical cupin fold and symmetric hexameric arrangement previously observed in homologous 11S globulins from other plant species^6,26,30,38–45^.

**Fig. 1:**
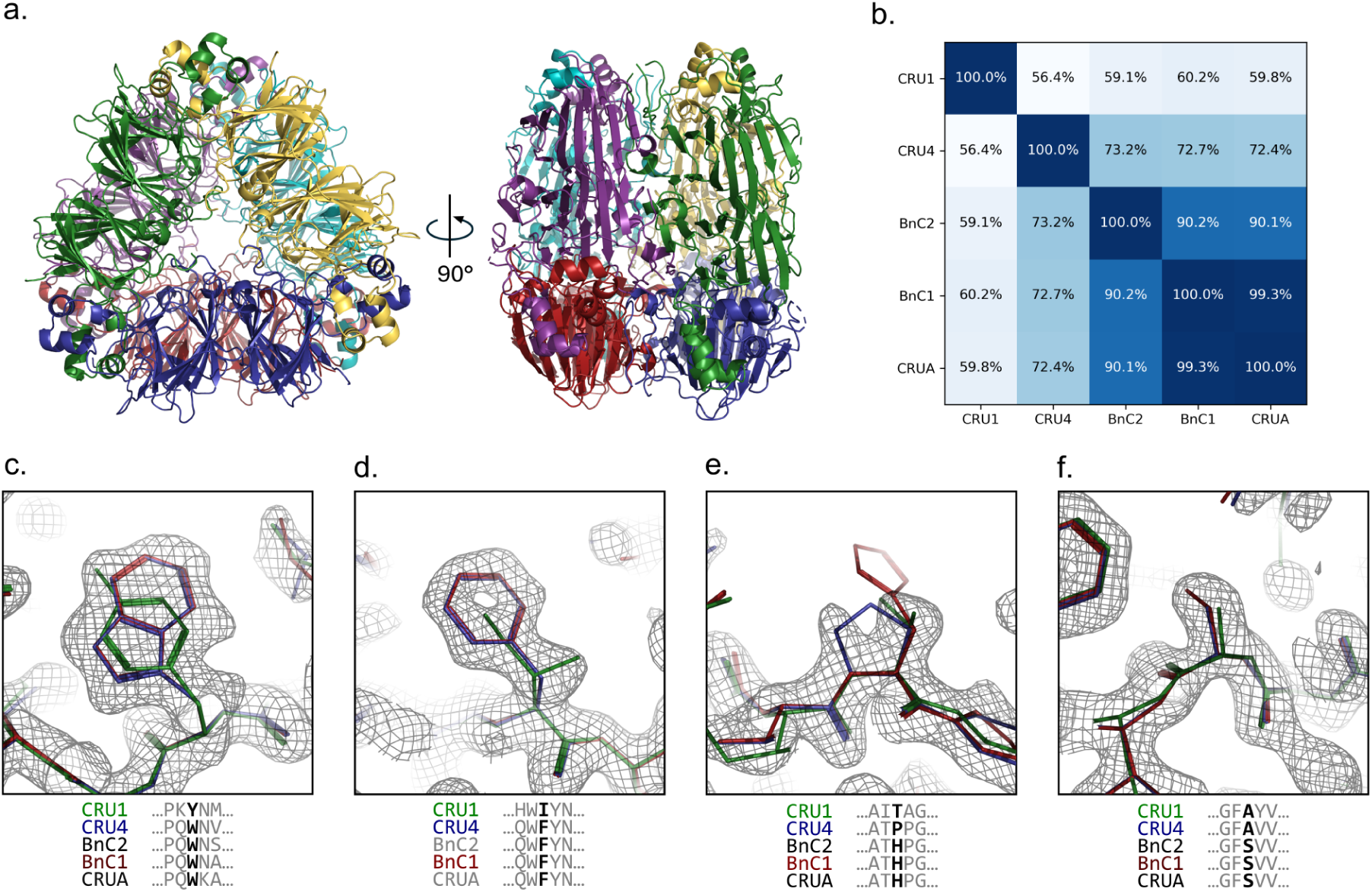
The hexameric reconstruction shows signs of heterogeneity. **a.** Cartoon representations of the refined homohexameric structure of the CRU4 isoform. Each of the six monomeric subunits are coloured distinctly. **b.** Heatmap showing the pairwise sequence identity between five annotated cruciferin isoforms. **c-f.** Details of the three refined structures of homohexameric cruciferin in stick representation overlaid alongside the density map. The three structures are drawn in different colours with, CRU1: green, BnC1: red, and CRU4: blue. Sections of a multiple sequence alignment (MSA) of all five reviewed cruciferin isoforms are shown underneath each render. **c.** Centered on Tyr360 of CRU1 / Trp341 of BnC1 / Trp318 of CRU4. **d.** Centered on Ile176 of CRU1 / Phe165 of BnC1 / Phe139 of CRU4. **e.** Centered on Thr169 of CRU1 / His158 of BnC1 / Pro132 of CRU4. **f.** Centered on Ala405 of CRU1 / Ser386 of BnC1 / Ala363 of CRU4. The same contour level is used for drawing the map in all four panels.

**Table 1:** Structural validation metrics for the homohexameric structures of the BnC1, CRU1, and CRU4 isoforms of cruciferin, all refined against the cryo-EM map at 1.8 Å resolution.

| **Isoform**<br>PDB code | **BnC1**<br>32LJ | **CRU1**<br>32LK | **CRU4**<br>32JP |
| --- | --- | --- | --- |
| **Composition** |  |  |  |
| Chains | 6 | 6 | 6 |
| Atoms | 18,616 | 18,458 | 18,555 |
| Residues (Protein) | 2,304 | 2,298 | 2,310 |
| Water Molecules | 478 | 446 | 477 |
| **Geometry** |  |  |  |
| MolProbity Score | 1.39 | 1.57 | 1.43 |
| Clash Score | 4.92 | 7.52 | 5.37 |
| Rotamer Outliers (%) | 1.51 | 1.61 | 1.52 |
| Cis Prolines (%) | 0.0 | 0.0 | 0.0 |
| Rama-Z Score (whole) | 0.34 | 0.72 | 0.33 |
| CaBLAM Outliers (%) | 0.86 | 1.04 | 0.59 |
| **Ramachandran Plot (%)** |  |  |  |
| Favored | 98.40 | 98.04 | 98.63 |
| Allowed | 1.60 | 1.96 | 1.37 |
| Outliers | 0.00 | 0.00 | 0.00 |
| **B-Factors (Mean)** |  |  |  |
| Protein | 20.85 | 21.48 | 21.19 |
| Water | 24.43 | 25.02 | 24.78 |
| **Model vs. Data** |  |  |  |
| Resolution (Supplied) | 1.8 Å | 1.8 Å | 1.8 Å |
| CC (mask) | 0.88 | 0.87 | 0.88 |

The pairwise sequence identities between five previously annotated isoform sequences are illustrated on Fig. 1b, spanning a broad range of sequence identities. Several regions of the reconstructed *D*_3_-symmetric map indicate local heterogeneity, reflecting the compositional diversity of the cruciferin sample. Instead of representing a single sequence, the density contains mixed contributions at positions where the known isoform sequences diverge (Figs. 1c-f). For instance, the map simultaneously accommodates multiple distinct sidechains such as overlapping signals from bulky tryptophan and tyrosine residues (Fig. 1c), phenylalanine and isoleucine residues (Fig. 1d), or proline and threonine residues (Fig. 1e). Overall, the consensus map exhibits a discernible bias toward the CRU1 and CRU4 isoforms, containing fewer features uniquely attributable to the three closely related isoforms CRUA, BnC1, and BnC2 (Fig. 1f illustrates a notable exception where density is uniquely attributable to the serine residue present in CRUA, BnC1, and BnC2).

To verify the isoform composition of the cruciferin sample it was analyzed by a peptide-based LC-MS approach, distinguishing and quantifying four groups of isoforms, as previously presented^28^. The resulting composition of this analysis is summarized in Supplementary Table 2, showing that three of the groups account for 98.7% of the cruciferin content of the sample. The most abundant isoforms are closely related to the reviewed CRU4 isoform at 48%, followed by the CRU1 isoform at 31%. One group, encompassing the three reviewed isoform sequences CRUA, BnC1, BnC2 together accounts for just 19%. The relative abundance of CRU1 and CRU4 correlates with the features observed in the *D*_3_ averaged cryo-EM map, where the density appears biased towards these isoforms.

### Distinction of isoforms by 3D classification

To resolve discrete compositional heterogeneity at the monomer level, approximately 1.8 million symmetry-expanded cruciferin monomer particles were processed by a sequential 3D classification approach, ultimately classifying them into three distinct classes with corresponding density maps built *de novo*. Density maps were reconstructed both from the full stack of 1.8 million particles at a resolution of 1.81 Å (the consensus map) and from the particles of each class at resolutions of 1.92, 1.96, and 1.93 Å. FSC and Guinier plots for the four reconstructions are shown in Supplementary Fig. 2.

Comparison of the consensus map to the three resulting class maps demonstrates clear resolution of the previously mixed local densities (Figs. 2a-e). While the consensus map ambiguously accommodates multiple overlapping sidechains at multiple positions, the isolated class maps show features matching specific isoforms. All highlighted differences between the maps demonstrate an enrichment of CRU1-like monomers in Class 1, BnC1-like monomers in Class 2, and CRU4-like monomers in Class 3. Additionally, we systematically quantified the agreement between each 3D class and 19 known sequences of 11S seed storage proteins from *B. napus* by comparing structures built using ModelAngelo^46^ against structures predicted using AlphaFold 3^47^. Unlike the bioinformatics tools implemented in ModelAngelo for identification of unknown proteins, this approach incorporates additional structural information. The results of the comparisons between the three classes and the 19 sequences are summarized in Fig. 2f as per-residue averaged log probabilities for every combination of class and sequence. While the consensus map has a similar degree of agreement with a range of diverse monomer sequences, each of the classes has a higher degree of similarity to a distinct subset of those sequences. By comparison to a phylogenetic tree of the sequences (Fig. 2f), it is apparent that the average log probabilities obtained for each class correlate with sequence similarity, and that the sets of sequences that match best with each class correspond to well-defined branches of the tree. The branches are consistent with the observations shown in Figs. 2a-e. The signs of isoform distinction between classes are distributed throughout the sequences, as illustrated in Supplementary Figs. 5-7, reporting ModelAngelo scores for each residue of CRU1, BnC1, and CRU4 against each of the three maps. Notably, these branches coincide with three out of four groups of sequences that can be identified by the peptide-based LC-MS quantification scheme. One remote branch of five sequences displays poor resemblance to any of the three classes, and we note that all evidence for these proteins has been at the genome level^48^.

**Fig. 2:**
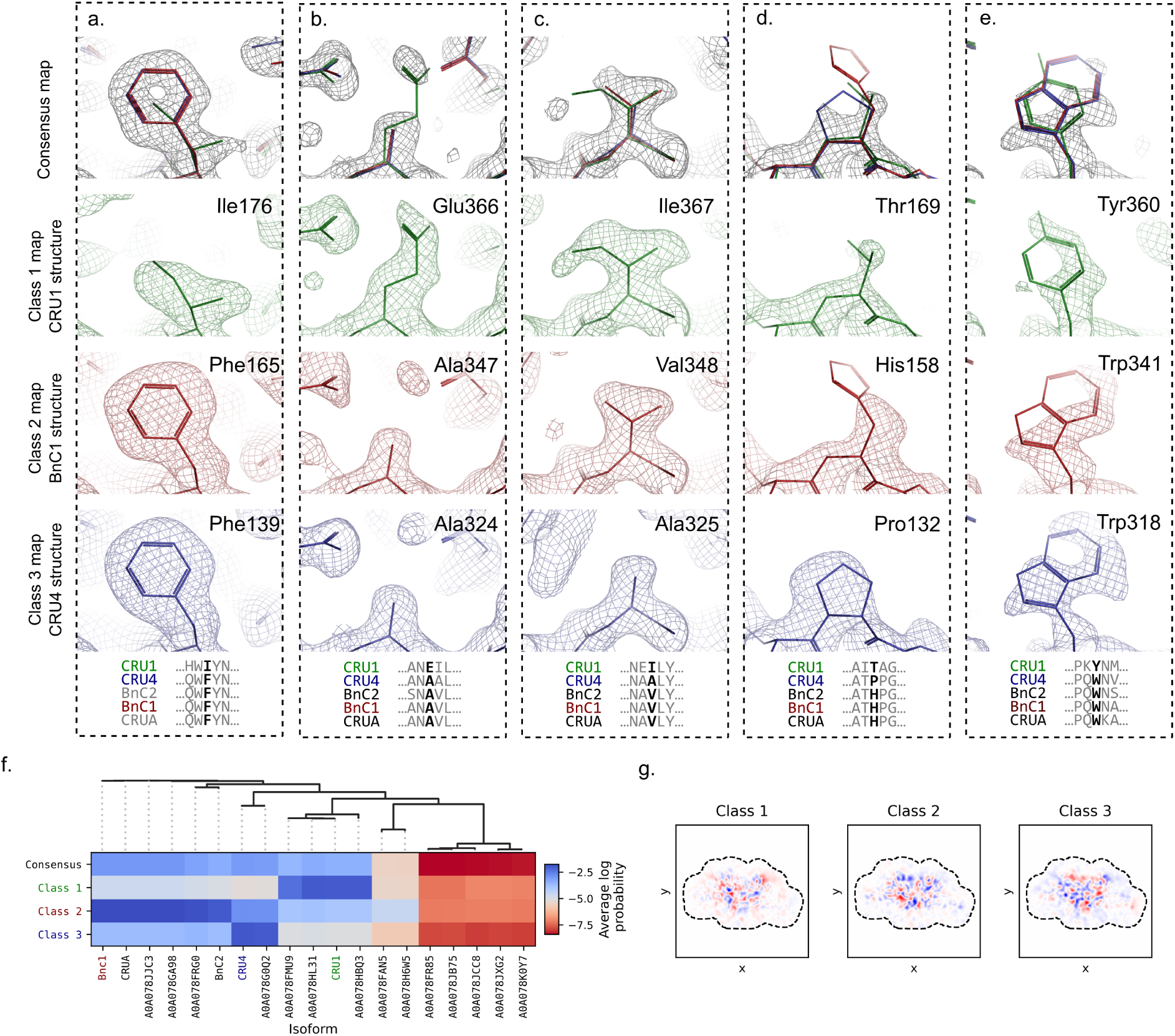
Monomer-level 3D classification distinguishes three groups of cruciferin isoforms. **a-e.** Examples of local differences between the densities of the consensus map and the three 3D class maps. The same five sites are shown for all four maps. In each panel, the consensus map reconstructed to a resolution of 1.81 Å is placed on top and shown in black mesh along with refined structures of three isoforms (green: CRU1, red: BnC1, and blue: CRU4) shown in stick representation. The maps of the three classes; class 1 (green, 1.92 Å), class 2 (red, 1.96 Å), and class 3 (blue, 1.93 Å) are shown alongside the same three structures. **a.** Centered on Ile176 of CRU1 / Phe165 of BnC1 / Phe139 of CRU4. **b.** Centered on Glu366 of CRU1 / Ala347 of BnC1 / Ala324 of CRU4. **c.** Centered on Ile367 of CRU1 / Val348 of BnC1 / Ala325 of CRU4. **d.** Centered on Thr169 of CRU1 / His158 of BnC1 / Pro132 of CRU4. **e.** Centered on Tyr360 of CRU1 / Trp341 of BnC1 / Trp318 of CRU4. **f.** Results from the comparison between ModelAngelo^46^ predictions for the sequences of both the consensus map and the densities of the three final 3D classes, and 19 known rapeseed cruciferin sequences. A heatmap shows the log probability for a match between each class and sequence, averaged per number of residues considered. A phylogenetic tree illustrates the relative similarities between the sequences. **g.** Projections of real-space difference maps of the three final density maps obtained by 3D classification. The heatmaps show projections of the difference maps by summation along the Z-axis, with blue denoting a positive sum of differences and red denoting a negative sum of differences. For projections along each of the Cartesian axes, see Supplementary Fig. 3. Grey dashed lines illustrate the extent of the boundaries of the focus mask used during 3D classification.

Real-space difference maps (Fig. 2g; Supplementary Fig. 3) reveal that the density variations are all found in the protein core. This confirms that distinguishing features originate from single monomers, rather than subtraction artifacts from adjacent subunits. Furthermore, these differences are distributed throughout the core rather than being localized to any particular region, indicating that numerous structural features enable the distinction between classes. This demonstrates that the classification approach is capable of distinguishing ∼50 kDa proteins with high sequence similarity and near identical folds.

### Misclassification rates and isoform quantification

The consecutive 3D classification approach appears to distinguish between isoforms at the single particle level. However, because cryo-EM datasets of small, highly similar proteins inherently exhibit low signal-to-noise ratios, separating these discrete states inevitably introduces a degree of classification ambiguity in the form of misclassifications^49^. To quantify the error rates, we generated simulated particles of pure isoforms, processed them using the classifier trained on the real cryo-EM data, and evaluated how often wrong class labels were assigned. Simulated particles with white noise represent an idealized scenario lacking the complex background noise and conformational diversity of real cryo-EM data. We assume that these factors are negligible relative to experimental noise and that simulated data adequately approximates experimental cryo-EM data.

The probabilities obtained from classification of simulated particles are summarized as a normalized confusion matrix in Fig. 3a, giving each of the probabilities *p_i,j_* of classifying any true particle belonging to class *i* to class *j*. Notably, correct classification of members of classes 1 and 2 is highly probable with *p*_1,1_ = 87.5% and *p*_2,2_ = 82.7%, while correct classification of members of class 3 is significantly less probable with *p*_3,3_ = 62.3%. One type of misclassification in particular, with probability *p*_3,2_ = 26.7%, is the largest source of error, indicating a notable blind-spot of the classifier. The overall distribution of the orientations of the monomeric particles is consistent with the *D*_3_ symmetry of the hexameric particles. As is apparent from Fig. 3c-d, the orientations of correctly and incorrectly classified particles do not perfectly follow the same distribution. Rather, the misclassification rate is dependent on the particle pose^49^. This can be understood, if projections of the distinct classes are more readily distinguishable from some orientations rather than others.

**Fig. 3:**
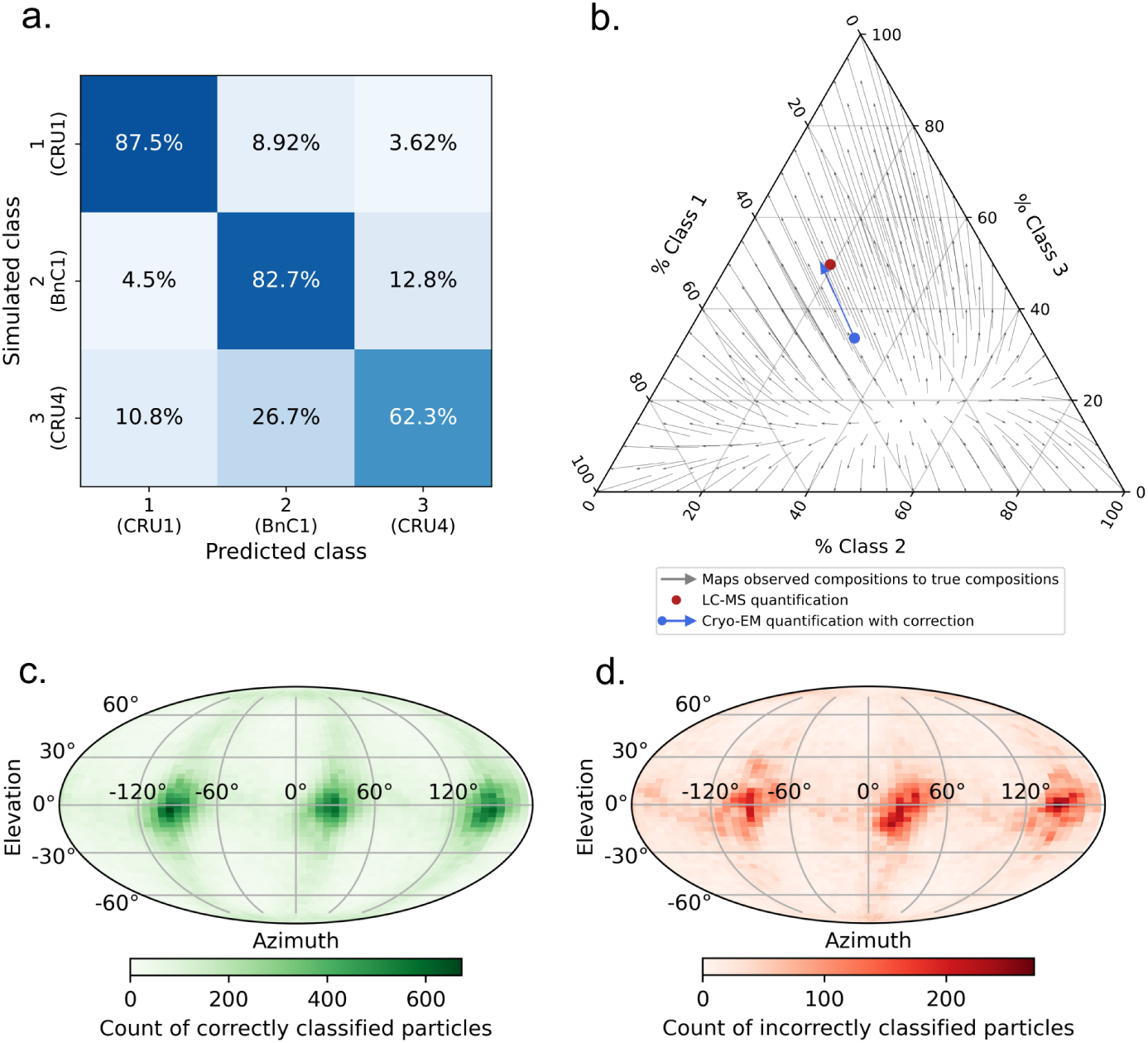
Assessment of classification accuracy and validation of overall isoform composition. **a.** Normalised confusion matrix for the classification of simulated particles of CRU4, CRU1 and BnC1, using the 3D classifier trained on the real particles. **b.** Ternary plot illustrating the apparent isoform class composition of the cruciferin sample. Arrows indicate the effect of the apparent misclassification rates, mapping observed compositions to true compositions. The red circle shows the composition obtained from quantification by LC-MS. The end of the blue arrow indicates the composition obtained from the 3D classification approach taking misclassification into account. **c-d.** Heatmaps of observed particle orientations of the simulated monomeric particles that were either correctly (**c.**) or incorrectly (**d.**) classified. The count of pairs of azimuth and elevation angles are plotted on a spherical surface using the Mollweide projection^50^.

Applying the estimated misclassification rates allowed us to correlate the apparent cryo-EM classification with the actual isoform composition. This yielded excellent agreement with the orthogonal LC-MS quantification (Fig. 3b), thus validating the 3D classification approach. The mass spectrometric quantification also shows evidence for a small fraction (1.3%) of cruciferin isoforms containing the pentapeptide sequence LEDFR (Supplementary Table 2), but none of the 3D classes showed notable similarity to either of the two known sequences containing this peptide (A0A078FAN5 and A0A078H6W5) (Fig. 2f). This small fraction has therefore been disregarded for purposes of comparison. Class 3, which is found to be the most populated at 48-50% of the monomers by the two methods, is also the least ambiguous at the sequence level, showing high similarity to just two known 11S globulins from rapeseed (with a mutual sequence identity of 99%). Despite this distinctness, it is also the class for which we observe the lowest fraction of correct classifications, illustrating that high prevalence is not the only determinant of effective distinction in 3D classification. The excellent agreement between the two independent estimates (from LC-MS and cryo-EM) of the overall subunit composition further supports the efficacy of 3D classifications for distinguishing cruciferin subunits at the monomer level and validates a quantitative interpretation of the results.

### Global thermodynamic modeling extracts interaction energies from population distributions

Having established that the classification workflow can reliably assign single-subunit signals from intact complexes, we next evaluate the resulting combinatorial space of the possible hexameric assemblies. There are 138 distinct ways to arrange combinations of three elements in *D*_3_ symmetry, each representing a possible arrangement of hexameric cruciferin built from three distinguishable subunits. Fig. 4a illustrates the apparent experimental frequency of each of these, plotted against the expected frequency based on total class populations, assuming no specificity in the interactions between subunits of different classes (the null model). All 138 possible arrangements are observed, although for some arrangements the frequency diverges significantly from the behavior expected from purely random assembly, showing either a lower or higher relative abundance. A selection of arrangements is illustrated in inset panels, while representations of all distinct hexameric complexes and their relative abundance can be found in Supplementary Fig. 8. Notably, the hexamers with the highest relative frequencies share a specific subarrangement of two subunits each of class 1 and class 3, indicating that this is a relatively favorable arrangement. Similarly, all 11 arrangements sharing a motif of three subunits labeled as class 1 arranged with three-fold rotational symmetry are among the arrangements with the lowest relative frequencies, indicating that this subarrangement is especially disfavored.

**Fig. 4:**
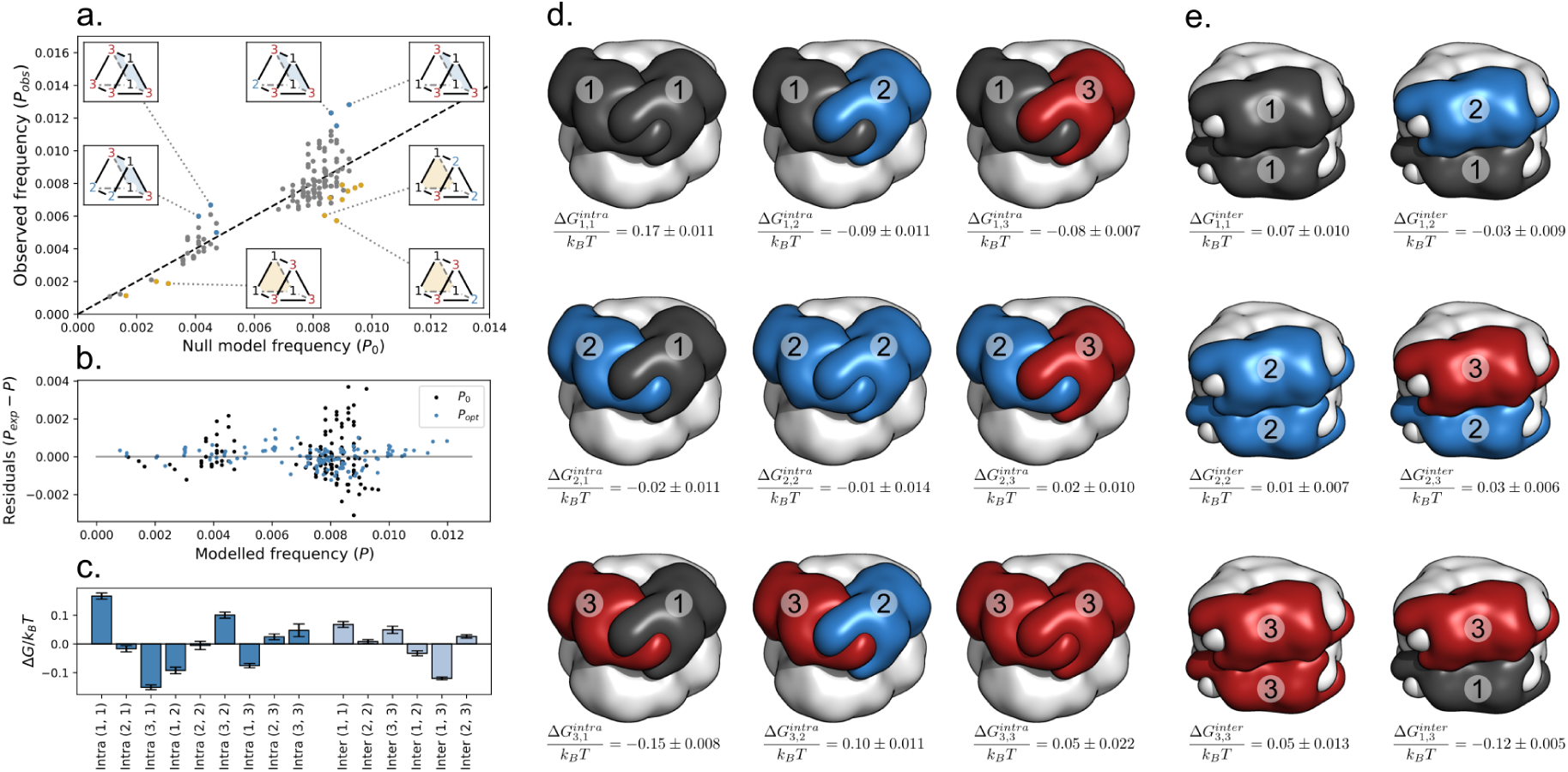
Rermodynamic modeling and estimation of interaction energies from the hexamer population. **a.** The apparent relative abundance of each of the 138 different possible arrangements of the three classes in a hexameric assembly of cruciferin with *D*_3_ symmetry. The black interrupted line illustrates the diagonal. Points above the diagonal indicate arrangements that are observed more often than expected from random assembly. Conversely, points below the diagonal indicate arrangements that are observed less often than expected from random assembly. Six arrangements with a shared subarrangement of two subunits each from class 1 and 3 are plotted in blue, with four such arrangements illustrated in inset panels with the shared subarrangement indicated by a blue plane. Another 11 arrangements with a shared subarrangement of three subunits of class 1 are plotted in yellow, with three such arrangements illustrated in inset panels with the shared subarrangement indicated by a yellow plane. All arrangements are represented in Supplementary Fig. 8, alongside their relative abundance. **b.** Residuals between the experimentally estimated frequency of each hexamer and frequencies modeled by the null model (*P*_0_, assuming no specificity between monomers) (in black) and the fitted model (*P_opt_*) (in blue). **c.** Estimated values of the 15 interaction energy fitting parameters from the optimal solution to the binding model. **d-e.** Graphical representations of the different possible intra-trimer (**d.**) and inter-trimer (**e.**) interactions between monomers of the three classes, along with the estimated parameter values from the optimal solution to the binding model. In **c-e.** the indicated errors are standard deviations estimated from the inverse of the Hessian matrix at the optimal solution.

Assuming equilibrium between all assemblies, a binding model describing the populations of all possible hexamers as a function of 15 interaction energies was fitted to the experimental population. In the model, the energy of each hexameric assembly is modeled as the sum of six intra-trimer interaction energies between adjacent pairs of monomers, and three inter-trimer interaction energies between adjacent pairs of monomers. Fig. 4b compares the experimental hexamer frequencies (*P_exp_*) with the frequencies modeled by the optimal, fitted model (*P_opt_*) and the null model assuming nonspecific interaction between classes of isoforms (*P*_0_). The null model shows significantly higher deviations from the experimental frequencies than the fitted model, especially at higher modeled population. The optimal values for the 15 interaction free energies all lie within *±*0.17*k_B_T* of their average, indicating subtle preferences in both intra- and inter-trimer interactions (Fig. 4c). All 15 interactions are illustrated on Figs. 4d-e along with the optimal fitted values. Intra-trimer interactions between two monomers of class 1 are the most unfavorable with Δ*G^intra^* = 0.17*k_B_T*, while both intra- and inter-trimer interactions between monomers of class 1 and 3 are relatively favorable with Δ*G^intra^* = *−*0.15*k_B_T* and Δ*G^inter^* = *−*0.12*k_B_T*, which is consistent with the preliminary observations illustrated in Fig. 4a. The average of all intra-trimer interaction energies and of all inter-trimer interaction energies are both fixed to 0, and the model thus does not report on the relative scale of the two groups of interaction free energies. Each of the 15 interaction types were also evaluated using two computational predictors of interaction energies^51,52^. However, no correlation to our estimated values was observed (Supplementary Fig. 10). We note that the sub-*k_B_T* differences between the estimated interaction energies are an order of magnitude smaller than reported expected errors of computational measures of interaction energies^51,53^. By random sampling of well-mixed populations and subsequent least-squares fitting, we estimate the detected free energies to be well above the detection floor of the method (Supplementary Fig. 9).

These observations all point to low but detectable levels of specificity in the relative arrangement of cruciferin isoforms. Still, the emerging picture is of a complex ensemble of distinct hexameric cruciferin assemblies coexisting, with a tremendous degree of promiscuity in the interactions between subunits. This might be advantageous for the plant as it allows for diversity in the expressed subunits, without hindering effective packing of proteins in the protein storage vacuoles. Conversely, the determined heterohexameric structure of an allergenic 11S globulin from hazelnut showed a specific arrangement of the individual subunits, representing a completely different assembly behavior^6^. The challenge of obtaining crystals of plant-derived soybean glycinin was recognized and attributed to subunit heterogeneity, and interestingly, glycinin from a mutant soybean expressing only a single subunit was crystallized and enabled the determination of the structure^30^. This case illustrates the exceptional strength of the present cryo-EM SPA approach for structural characterization of heterogeneous biomolecular complexes from endogenous sources that are fundamentally incompatible with crystallization.

## Conclusions

This work establishes structural thermodynamics as a method based on cryo-EM to resolve and quantify compositional heterogeneity in biomolecular assemblies of structurally similar protein isoforms. By applying iterative 3D classifications to native *B. napus* cruciferin, we distinguished three groups of highly similar isoforms within hexameric assemblies. The efficacy of the method was validated through classification of simulated particles and corroborated by orthogonal LC-MS measurements in agreement with the composition derived from cryo-EM. The method allows for a detailed and quantitative description of the interactions between distinguishable subunits, and resolves sub-*k_B_T* energetic multimerization preferences.

The methodology presented could be applied to elucidate the compositional heterogeneity of diverse multimeric protein systems that may display promiscuous interactions between subunits. Such systems are fundamentally challenging to characterize in bulk, necessitating single-particle approaches. By illustrating the feasibility of 3D classifications at the single subunit level for the characterization of compositional heterogeneity, we provide a reliable framework for the structural and energetic study of heterogeneous systems with broad implications for health and disease. Notable examples are the human GABA_A_ receptor^4,54–57^ and the 20S proteasome^58,59^ where promiscuous or interchangeable subunits complicate bulk characterization.

To apply structural thermodynamics to other heterogeneous assemblies, its success will rely on the intrinsic interplay between subunit similarity, achievable resolution, and classification accuracy. Distinguishing highly similar subunits inherently demands high resolution densities; if the resolution is insufficient to capture structural differences between subunits, the misclassification rates will inevitably increase and hinder accurate quantitative interpretation. This relationship is cyclical, as elevated misclassification rates obscure the populations of each class, which in turn lowers the achievable resolution of the maps reconstructed from the isolated classes. This method demonstrates a thermodynamic interpretation of cryo-EM data, enabling a comprehensive understanding of assembly diversity of very similar subunits in native biomolecular complexes. Cruciferin serves as a challenging case study in subtle assembly preferences, but biomolecular systems with higher subunit specificity are equally amenable to this framework.

## Materials and methods

### Cryo-EM grid preparation and data collection

Purified cruciferin derived from cold-pressed rapeseed and further purified as previously described^60^ was kindly provided by Taco Nicolai at Le Mans Université. The lyophilized powder was dissolved in 10 mM Tris-HCl pH 8.5, 500 mM NaCl, to a final protein concentration of 0.96 mg/ml, based on a cruciferin concentration of 80% w/w in the powder^60^. Copper 300 grids with Quantifoil R 1.2/1.3 mesh were glow discharged for 45 s at 15 mA and loaded in a Vitrobot Mark IV (Thermo Fischer Scientific) equilibrated at 4 *^◦^*C and 100% humidity. Three microliter of the cruciferin solution was transferred to the mesh, which was promptly blotted for 3.5 s before plunging into liquid ethane.

Data was collected on a Titan Krios G3i (Thermo Fisher Scientific) with a K3 detector, using a magnification of 130,000x and a resulting pixel size of 0.647 Å. EPU software (Thermo Fisher Scientific) was used for automated data collection, capturing a total of 6781 exposures at targeted defocus values at steps of 0.2 µm between −0.3 µm and −1.5 µm. A total exposure dose of 59.5 e/Å^2^ was applied.

### Cryo-EM data processing and model building

All cryo-EM data processing was performed using CryoSparc^20^. Following initial patch motion correction and CTF (contrast transfer function) estimation, manual curation of the exposures resulted in 3414 accepted exposures. Blob picking on a subset of 100 denoised micrographs followed by a round of 2D classification and *ab initio* reconstruction with threefold dihedral symmetry enforced (*D*_3_ in Schoenflies notation) gave an initial volume that was used for template picking. Template picking, another round of 2D classification, and heterogeneous refinement gave a stack of 309,000 particles. With local and global CTF refinements (including fitting of tilt, trefoil, spherical aberration, tetrafoil, and anisotropic magnification) as well as reference-based motion correction, a reconstruction of hexameric cruciferin with an overall resolution at 1.83 Å was achieved as evaluated by the FSC curve at a value of 0.143.

Models of homohexameric rapeseed cruciferin were built using initial structures predicted using AlphaFold^47^ with an input of six copies each of the sequences of three different cruciferin isoforms with gene names CRU1 (UniProt: P33525), BnC1 (UniProt: P33523), and CRU4 (UniProt: P33522). The predicted structures were refined by real-space refinement to the sharpened density maps using Phenix^61^ and Coot^62^. Molecular figures were prepared and rendered using PyMOL^63^.

### 3D classification of monomeric subunits

For 3D classification of cruciferin at the monomer level, a stack of all monomeric subunits was prepared by symmetry expansion and subtraction of the estimated signal of the remaining subunits. The stack of hexameric particles were expanded according to their *D*_3_ symmetry using the symmetry expansion method in CryoSparc. For particle subtraction, a mask encompassing five of six monomers of the determined hexameric structures was used. This mask was prepared by first using the ‘molmap’ command in UCSF ChimeraX (version 1.6.1)^64^, with a selection of all atoms of chains B-F of the refined structure of the homohexamer of the CRU4 isoform and a resolution parameter of 12 Å. A similar volume around chain A was also prepared, and subtracted from the first. The resulting volume was turned into a mask in CryoSparc, using a lowpass filter of 20 Å and a soft padding width of 12 pixels. The mask was used for subtracting the expected signal from the five subunits adjacent to each subunit, using the ‘particle subtraction’ method in CryoSparc.

The resulting particles acted as the input of a series of consecutive 3D classifications, by an iterative approach searching for incrementally better apparent separation of isoforms (as evaluated by the identification approach outlined in the next section). A focus mask for 3D classifications of the monomeric subunits was prepared by the same approach as for the mask for particle subtraction, but this time taking the volume around chain A of the refined structure of homohexameric CRU4 isoform cruciferin, and subtracting the volume around the five remaining chains. This volume was turned into a mask in CryoSparc, again using a lowpass filter of 20 Å and a soft padding width of 12 pixels. This mask was used for all 3D classifications. The apparent optimal results were reached by performing an initial unsupervised 3D classification in CryoSparc of all particles into five classes at a target resolution of 3 Å, with a fixed Online Expectation Maximisation (O-EM) learning rate of 1.0 for five O-EM epochs, default convergence criteria thresholds, and forcing hard classification. The resulting density maps from the first 3D classification were used to initialise a second 3D classification using an initial low-pass filter at 2 Å, this time with the learning rate set to 0.1 and without forcing hard classification. Maps and particles from the second classification were aligned using the ‘align 3D maps’ method in CryoSparc, and used as input for a third classification with settings identical to the second. The five resulting classes were regrouped into three superclasses using the ‘regroup 3D classes’ method in CryoSparc. A final soft 3D classification was performed, initialised by the reconstructed densities of the three superclasses, again using an initial low-pass filter of 2 Å and a fixed O-EM learning rate of 0.1, this time with stricter convergence criteria of 0.001 for the density change threshold and 1% for the particle class change criterium. In all further analyses, only the most likely class label after the final classification is considered for each particle.

Experiments with 3D classification of smaller subsets of the stack of monomeric particles were attempted, and although the runs converged with much shorter runtimes, we observed a poorer apparent distinction of cruciferin isoforms. This observation can be understood as each class of particles must be populated enough to reconstruct a density map with high enough resolution relative to the distinctions aimed for. Additionally, we observed no apparent benefits in terms of improved separation of isoforms, by using a target resolution higher than 3.0 Å in 3D classifications.

### Isoform identification

To evaluate if 3D classifications can distinguish between monomeric cruciferin isoforms, we employed a method that combines automated model building using ModelAngelo^46^ with structural information from structures predicted by AlphaFold 3^47^. A total of 19 sequences annotated as 11S seed storage protein from *B. napus* in UniProt^32^ were selected, 5 of which are manually annotated and reviewed^34–37^, and 14 originate from sequencing of the *B. napus* genome^48^. A multiple sequence alignment of the sequences was performed using ClustalW, and a phylogenetic tree was drawn using toytree^65,66^. The structures of each protein was predicted using AlphaFold 3^47^ and superposed with a consensus map obtained from all 1.8 million monomeric particles using ChimeraX^64^. A total of 193 sets of *α*-carbon atoms were found within a threshold distance of 2 Å from each other across all superposed structures, and regarded as regions with near perfect structural conservation.

The structures and sequences of each 3D class and of the consensus group of all 1.8 million monomer particles were predicted from their refined density maps using ModelAngelo^46^ without providing any sequence information. At each residue recognised by ModelAngelo that coincides with one of the conserved *α*-carbons (being itself within a distance of 2 Å), the probability of a match with the residues found at the given position in each of the 19 sequences was obtained from the ModelAngelo hidden Markov model (HMM) profiles. For the consensus map, 152 residues were found at matching positions, while 138-145 matching residues were found for the three 3D classes. The average of all log probabilities of matches is used as a measure of how well a map reconstructed from a 3D class matches with a given sequence.

### Estimating misclassification rates by simulated particles

To approximate the error-rates of the 3D classification approach assuming that each of the three classes truly corresponds to a group of isoforms, simulated particles of pure isoforms were generated and classified using the classifier that was trained on the real particles. Structures of three monomeric cruciferin isoforms representative of apparent members of the three classes (CRU1 from class 1 (UniProt: P33525), BnC1 from class 2 (UniProt: P33523), and CRU4 from class 3 (UniProt: P33522)) in their mature cleaved forms were predicted using AlphaFold3^47^. To avoid any influence from dynamic regions of the protein, all residues in the predicted structures with low local confidence scores (pLDDT < 50) were removed, as this is a useful predicted measure of protein disorder^67^. Densities around each pruned structure were generated at a resolution of 2 Å using ChimeraX^64^, and used for simulating particles in CryoSparc^20^ using a signal-to-noise ratio of 0.15 and sampling CTF estimates and orientations from the real symmetry-expanded particles. For each of the three isoforms, 144,000 particles were simulated using different subsets of the real particles for sampling CTF estimates and orientations. The simulated particles were classified using the final classifier trained on the real particles. This was achieved by initializing a 3D classification with the final density maps of the real classification low-pass filtered at 2 Å, setting the O-EM learning rate to 0, the filter resolution to 3 Å, and terminating the classification after a single iteration of the Full-Batch Expectation Maximisation (F-EM) algorithm.

### Hexamer binding model

A binding model was devised to model the apparent populations of each of the 138 distinguishable hexameric cruciferin assemblies based on their isoform composition and arrangement. This approach assumes that the energy of each possible hexameric assembly can be calculated by summation of energetic contributions of interactions between pairs of adjacent monomers specific to the assembly. Hexameric cruciferin with *D*_3_ symmetry can be considered as a dimer of trimers. The energy of each of the possible hexamer arrangements is modelled as the sum of the energies of six interactions between adjacent pairs of monomers within the two trimers (intratrimer interactions) and three interactions between adjacent pairs of monomers between trimers (inter-trimer interactions). These interactions account for the bulk of the molecular contacts between monomers within the hexameric assemblies.

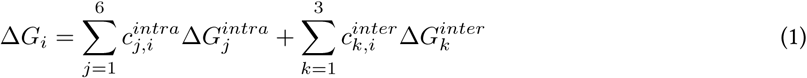

Where Δ*G_i_* represents the dimensionless energy (scaled by *k_B_T*) of hexamer *i*, and *c_j,i_* and *c_k,i_* are the integer counts of each interaction of type *j* or *k* within hexamer *i*. For hexamers composed of monomers from three distinguishable classes, there are nine distinct intra-trimer interactions and six distinct inter-trimer interactions to consider. The pairwise interaction energies are not absolute but rather defined relative to each other, by constraining the sum of all intra-trimer interaction energies and all inter-trimer interaction energies to be 0:

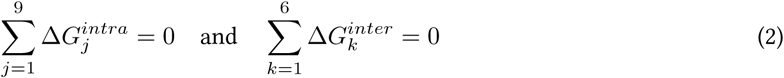

A statistical weight for each distinct hexameric assembly, *W_i_*, can be defined from its energy, the multiplicity of the hexamer arising from symmetry, *m_i_* (the number of distinct arrangements arising from applying the *D*_3_ symmetry operations to hexamer *i*), the monomer-composition of the hexamer (where *n*_1*,i*_ + *n*_2*,i*_ + *n*_3*,i*_ = 6), and the concentration of free monomers of each of the three classes ([*C*_1_], [*C*_2_], and [*C*_3_]). The concentration of each hexamer follows the following relation:

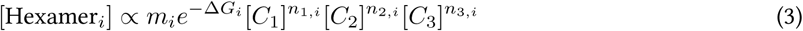

By rearrangements, the weights of each hexamer, *W_i_*, can be defined in terms of relative free monomer concentrations (i.e. [*C*_1_]/[*C*_3_] and [*C*_2_]/[*C*_3_]). Because all assemblies are hexameric (*n*_1*,i*_ + *n*_2*,i*_ + *n*_3*,i*_ = 6 for all *i*), [*C*_3_]^6^ can be factored out as a constant for all species.

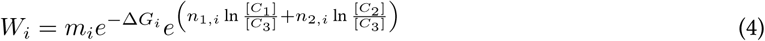

The relative populations of each hexameric species are then calculated as the ratios between the weights and the partition function, i.e. the sum of all weights.

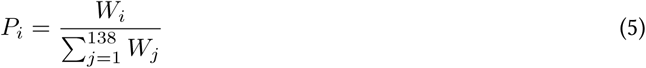

To account for the misclassification of particles, the populations are corrected for the apparent misclassification rates of each hexamer by constructing a 138 *×* 138 matrix **A** for which each element **A***_ij_* is the probability of misinterpreting hexamer *j* as hexamer *i*. This matrix is constructed from the confusion matrix for monomer classification as probed by classification of simulated particles (Fig. 3a). The corrected modelled population, *P ^′^*, is then:

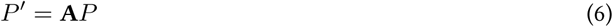

The relative concentrations of free monomers of each class were used as two additional free parameters, namely [*C*_1_]/[*C*_3_] and [*C*_2_]/[*C*_3_]. An L2 regularization penalty was applied to the interaction energy parameters with a regularization parameter of *λ* = 10*^−^*^4^, chosen based on the L-curve criterion^68^. The two parameters describing the relative concentrations of free monomers were not regularized. This resulted in the following objective function:

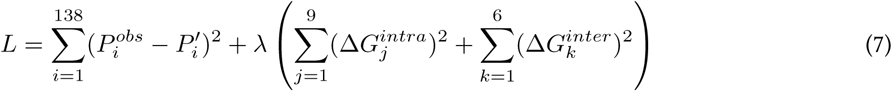

The objective function was minimized using the L-BFGS-B algorithm as implemented in the scipy.optimize package^69^. Standard deviations for the parameter estimates were calculated from the inverse of the Hessian matrix at the optimal solution.

### Computational prediction of interaction energies

Dimer structures representing every pair of inter- and intra trimer interactions between BnC1, CRU1, and CRU4 were prepared by extracting select monomers from the three refined hexameric structures. All structures were optimized using FoldX^52^ to eliminate clashes. Interaction energies between monomers were assessed using both FoldX and the PRODIGY webserver^51,70^.

### Peptide based LC-MS isoform quantification

Quantification of cruciferin isoforms was performed by a peptide-based LC-MS approach as previously described^28^. In brief, four groups of cruciferin variants were quantified by LC-MS/MS using selected reaction monitoring of peptides after digestion with trypsin: 1. includes the annotated isoform CRU1 (UniProt accession P33525), 2. includes the annotated isoforms CRUA, BnC1, and BnC2 (UniProt accession P11090, P33523, and P33524), 3. includes the annotated isoform CRU4 (UniProt accession P33522), and 4. includes two unannotated cruciferin variants with the UniProt accessions A0A078FAN5 and A0A078H6W5. These four groups were quantified using the target peptides VEHVR, VEHIR, VEHLR, and LEDFR, respectively. This is summarised in Table 2. A surrogate matrix-matched external calibration curve with synthetic peptide standards were used to quantify the percentage of each variant group of the total protein content in the sample measured by the Dumas method.

## Acknowledgements

The authors thank the Novo Nordisk Foundation for funding through their challenges program (NNF21OC0065495 SEEDFOOD). LC-MS data was generated by accessing LC-MS equipment at Department of Food Science, University of Copenhagen funded by FOODHAY (Food and Health Open Innovation Laboratory, Danish Roadmap for Research Infrastructure).

## Supplementary

### Extended structural descriptions

Several sites of the reconstructed *D*_3_ hexameric map show convincing evidence of local heterogeneity. At positions where the isoform sequences differ, the density shows clear contributions from different residues, as exemplified in Figures 1c-f where four local regions of the density are shown alongside the three refined homohexamer structures that cover the local sequence diversity of the isoforms. At one position, tryptophan residues are found in four out the five reviewed isoform sequences, while a tyrosine residue is found in the equivalent position of the CRU1 isoform (Fig. 1c). The density is able to accommodate the bulky tryptophan side-chain but also features additional density consistent with the hydroxy group of Tyr360 of CRU1. At another site where CRU1 features an isoleucine residue, while all other sequences feature a phenylalanine residue (Fig. 1d), density satisfying both Ile and Phe is present. Importantly, no other rotamer of Phe is feasible in this position due to steric clashes, and thus, a double conformation of Phe would not be able to explain the density accommodating the C*_γ_*_2_ atom of Ile176 of the refined structure of CRU1. At another position where CRU1 features a threonine residue, CRU4 features a proline residue, and the remaining three isoforms share a histidine (Fig. 1e). Here, the density accommodates Thr169 of CRU1 and also shows clear evidence for the tertiary amide of Pro132 of CRU4, with density that is explicable by the C*_δ_* atom of proline but not by any other residue. Several rotamer conformations are plausible for the His158 of BnC1, which could explain the lack of density. Overall, the reconstructed map shows a bias towards CRU1 and CRU4, as few sites in the map cannot be accounted for by the sequences of these two isoforms. One example is a site where the density fully accommodates the O*_γ_* atom of Ser386 of BnC1 (which is shared as a serine with CRUA and BnC2) while an alanine is found in the same position of both CRU1 or CRU4 (Fig. 1f). Based on these highlighted regions of the density map, evidence is found of contributions from three distinguishable groups of residues: CRU1; CRU4; and the three more closely related isoforms CRUA, BnC1, and BnC2.

Figures 2a-e illustrate examples of five regions of the densities of both the consensus map reconstructed from all monomeric particles and the three maps reconstructed after 3D classification, that each show signs of separation of isoforms. All highlighted differences between the maps are consistent with an enrichment of particles of cruciferin monomers similar to the CRU1 isoform in class 1, monomers similar to the BnC1 isoform in class 2, and monomers similar to the CRU4 isoform in class 3. Fig. 2a (which highlights the same region as Fig. 1d for the reconstruction of the hexameric assembly) shows that while the consensus map accommodates Ile176 of CRU1, Phe165 of Bnc1, and Phe139 of CRU4, the maps after classification only fully accommodate either an Ile or a Phe sidechain. Similarly, the map from class 1 shows additional density at the site of Glu366 of CRU1, compared to classes 2 and 3, which both show tighter densities consistent with the Ala residue present in all isoforms except CRU1 (Fig. 2b). At another site shown in Fig. 2c, the class 1 map accommodates the full sidechain of Ile367 of CRU1, while the class 2 map accommodates Val348 of BnC1 but does not have density to encompass the C*_δ_* atom of an Ile residue, and the map of class 3 accomodates Ala325 of CRU4 but does not explain the C*_γ_*_1_ and C*_γ_*_2_ atoms of either an Ile or Val residue. The consensus map shows density consistent with the C*_δ_* atom of Pro132 of CRU4 (Fig. 2d) at a position where only this isoform has a Pro residue. This feature, which is not explicable by any other residue, is also present in the map of class 3 but not present in the maps of the two other classes (Fig. 2d). While the sidechains are not completely resolved in the reconstructions of the classified particles, the heterogenic site near Tyr360 of CRU1 (which also showed signs of heterogeneity in the hexameric reconstruction (Fig. 1a)) show signs of isoform separation, as the class 1 map is consistent with Tyr360 of CRU1 and inconsistent with a Trp residue, while class 2 and 3 maps are consistent with Trp residues of BnC1 (Trp341) and CRU4 (Trp318) while being inconsistent with a Tyr residue (Fig. 2e).

The high-resolution structures of native cruciferin show no evidence of post-translational modification (PTMs). Although phosphorylation of specific residues was detected in cruciferin homologs from *A. thaliana*^71^, no unattributable densities were observed adjoining serine, threonine, or tyrosine residues in the refined structures of homohexameric cruciferin from *B. napus*. In a previously published crystal structure of recombinantly produced procruciferin, a ligand was observed at a site adjacent to a tryptophan and three tyrosine residues, which was attributed to glycerol^26^. No density was detected at the corresponding site in the cryo-EM reconstruction.

### Scaling behavior of thermodynamic modeling in symmetric assemblies

In a *D*_3_-symmetric hexameric system composed of *n* distinguishable subunits, the number of distinct hexameric arrangements, *N_D_*, scales as a sixth-order polynomial according to the Cauchy-Frobenius Lemma, by counting the total number of arrangements left unchanged by applying each symmetry operation, *g*, divided by the number of operations, *|G|*.

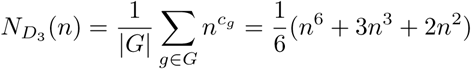

Where *c_g_* is the number of disjoint cycles of objects shuffled by the operation *g*. To clarify, all *n*^6^ possible arrangements are left unchanged by the identity operation. For each of the three rotations about the axes with twofold symmetry, there are 3 disjoint cycles, resulting in *n*^3^ unchanged arrangements per rotation. Finally, for each of the two rotations (120*^◦^* and 240*^◦^*) about the axis with threefold symmetry, there are 2 disjoint cycles, resulting in *n*^2^ unchanged arrangements per rotation. For the present cruciferin system where we can distinguish 3 groups of isoforms, there are *N_D_* (3) = 138 distinct hexameric assemblies. Similar expressions likewise exist for any other symmetry. Supplementary Table 1 gives counts of assemblies for values of *n* up to 6 for select symmetries in addition to *D*_3_ symmetry of hexameric cruciferin: *C*_3_ as the symmetry of vicillin-like globulin SSPs, *C*_5_ as the pseudosymmetry of the GABA_A_ receptor, and *D*_7_ as the symmetry of the GroEL chaperone.

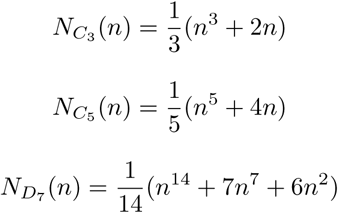

It is a common feature of assemblies of all symmetries, that the order of the polynomial describing *N* (*n*) equals the oligomeric state of the assembly. *N* can therefore scale drastically with increasing numbers of distinguishable subunits. Meanwhile, the number of free fitting parameters will only scale quadratically with increasing numbers of distinguishable subunits (for a non-staggered dihedral symmetry as presented for cruciferin, the bulk of contacts between subunits can be described by 2*n*^2^ *− n* pairwise interactions). This is an inherent strength of the global thermodynamic modeling approach in handling systems of more distinct subunits, as the number of datapoints available (*N*) scales much faster than the degrees of freedom of the model, ensuring overdetermination of the system.

**Supplementary Table 1:**
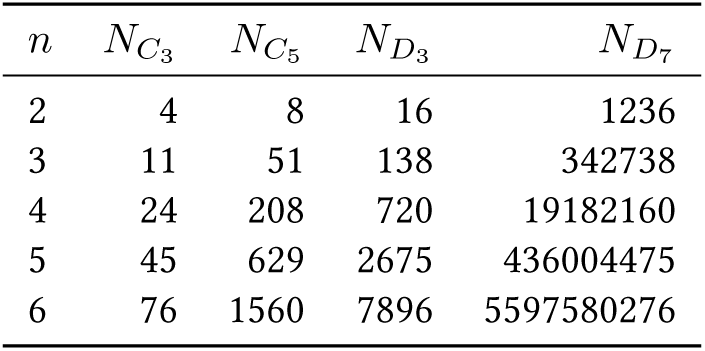
The number of possible arrangements of *n* different subunits for values of *n* between 2 and 6, for assemblies of each of the exemplified symmetries.

| $n$ | $N_{C_3}$ | $N_{C_5}$ | $N_{D_3}$ | $N_{D_7}$ |
| --- | --- | --- | --- | --- |
| 2 | 4 | 8 | 16 | 1236 |
| 3 | 11 | 51 | 138 | 342738 |
| 4 | 24 | 208 | 720 | 19182160 |
| 5 | 45 | 629 | 2675 | 436004475 |
| 6 | 76 | 1560 | 7896 | 5597580276 |

### Supplementary Figures

**Supplementary Table 2:** Overview of the isoform composition of the cruciferin sample, as determined by LC-MS, along with the isoforms associated with each peptide identified by their UniProt ID, entry name, and gene names. For each isoform, the preferred identifier used in this work (either UniProt ID or gene name) is shown in bold.

| Peptide | UniProt ID | UniProt Entry Name | UniProt Gene Name | Percentage (%) |
| --- | --- | --- | --- | --- |
| VEHIR | P33523 | CRU1_BRANA | **BnC1** | 19.1 |
|  | P11090 | CRUA_BRANA | **CRUA** |  |
|  | **A0A078JJC3** | A0A078JJC3_BRANA | BnaC07g48660D |  |
|  | **A0A078GA98** | A0A078GA98_BRANA | BnaA06g36310D |  |
|  | **A0A078FRG0** | A0A078FRG0_BRANA | BnaA02g22500D |  |
|  | P33524 | CRU2_BRANA | **BnC2** |  |
| VEHVR | P33525 | CRU3_BRANA | **CRU1** | 31.4 |
|  | **A0A078HBQ3** | A0A078HBQ3_BRANA | BnaA01g08350D |  |
|  | **A0A078FMU9** | A0A078FMU9_BRANA | BnaA08g13680D |  |
|  | **A0A078HL31** | A0A078HL31_BRANA | BnaC01g09900D |  |
| VEHLR | P33522 | CRU4_BRANA | **CRU4** | 48.3 |
|  | **A0A078G0Q2** | A0A078G0Q2_BRANA | BnaC05g02160D |  |
| LEDFR | **A0A078FAN5** | A0A078FAN5_BRANA | BnaC08g43570D | 1.3 |
|  | **A0A078H6W5** | A0A078H6W5_BRANA | BnaA09g50780D |  |

**Supplementary Fig. 1:**
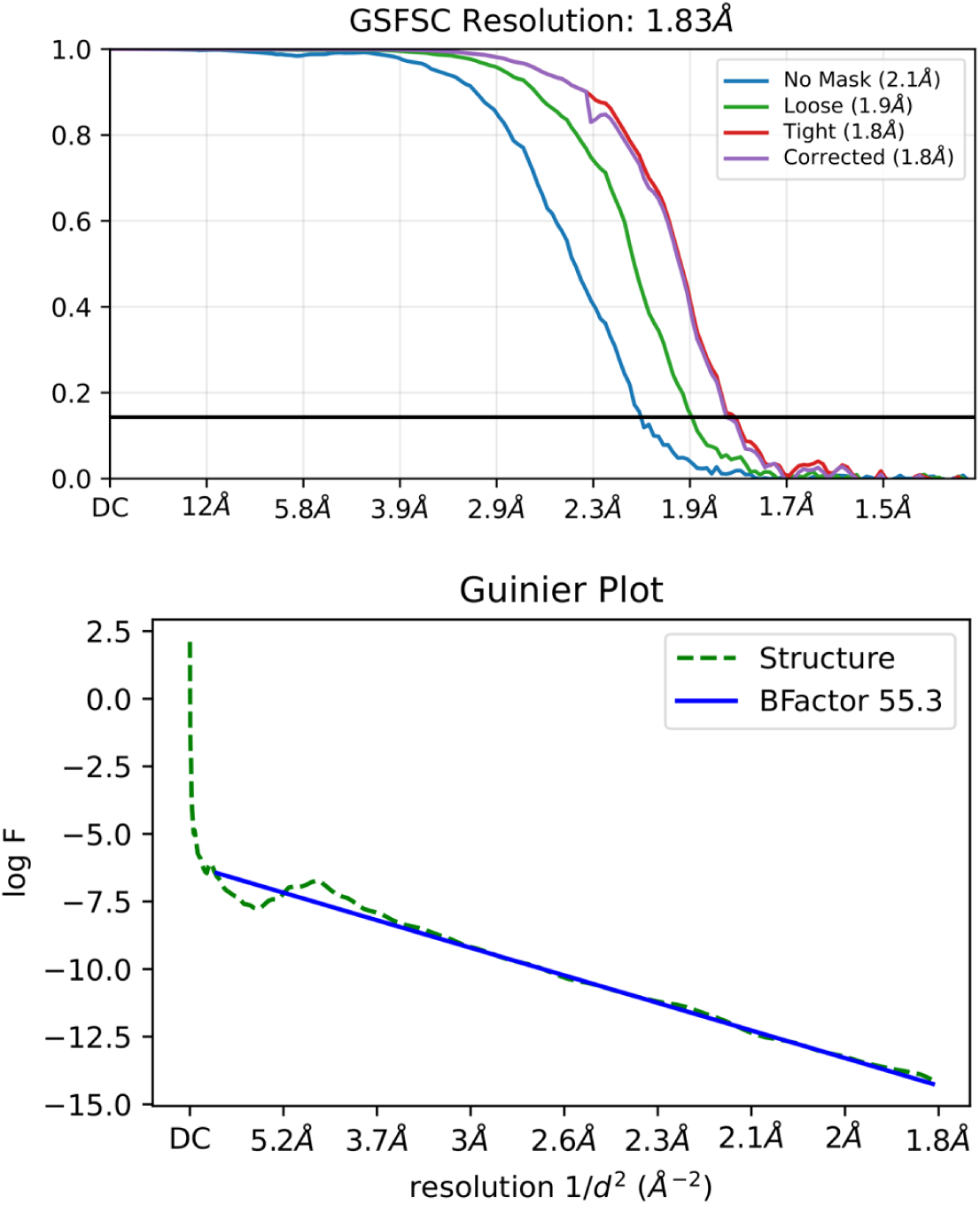
Gold standard FSC (Fourier Shell Correlation) and Guinier plots for the 3D reconstruction of hexameric cruciferin with *D*_3_ symmetry enforced.

**Supplementary Fig. 2:**
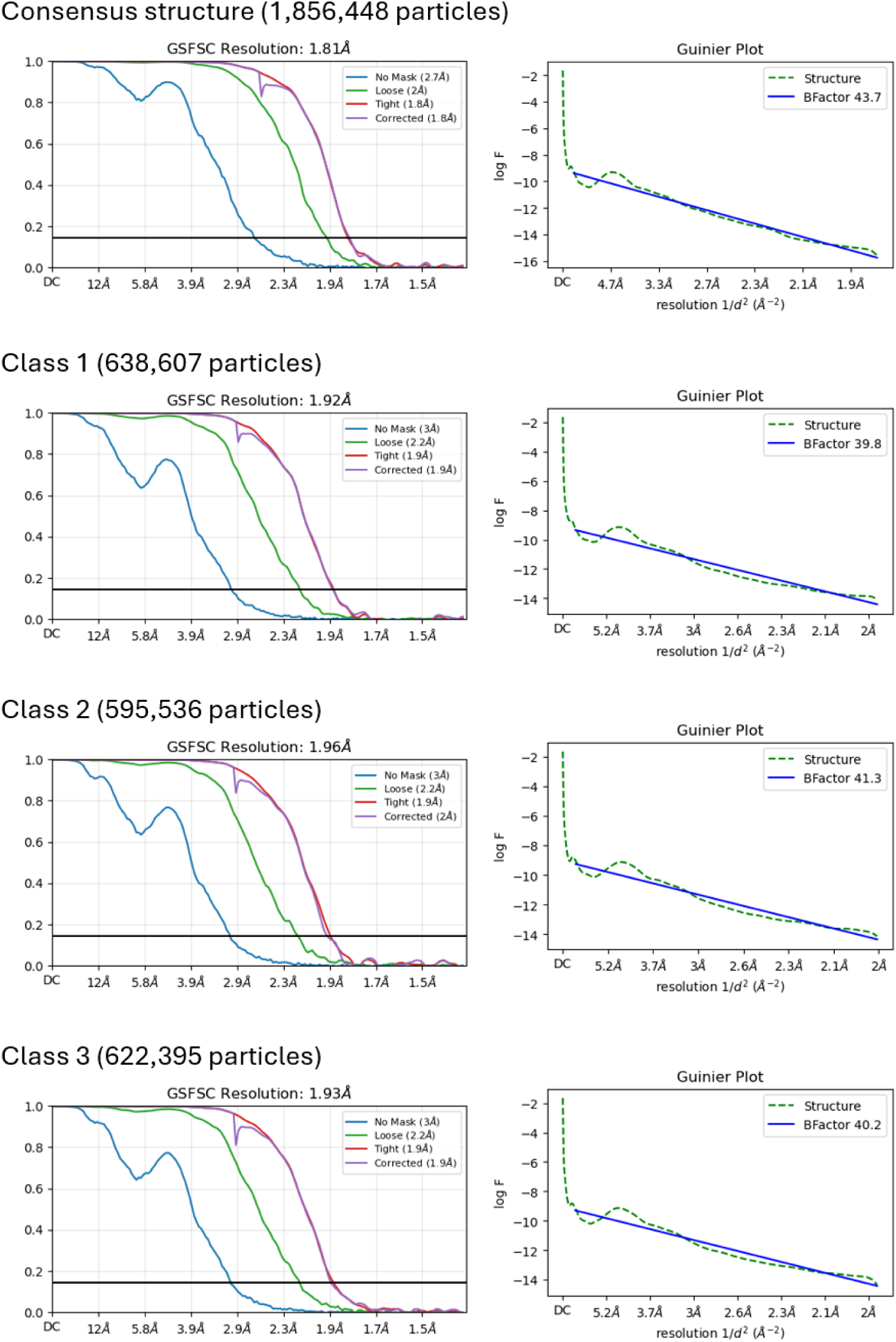
Gold standard FSC (Fourier Shell Correlation) and Guinier plots for the 3D reconstructions of the consensus volume from all monomeric particles after symmetry expansion and particle subtraction, and the particles of each of the three final 3D classes.

**Supplementary Fig. 3:**
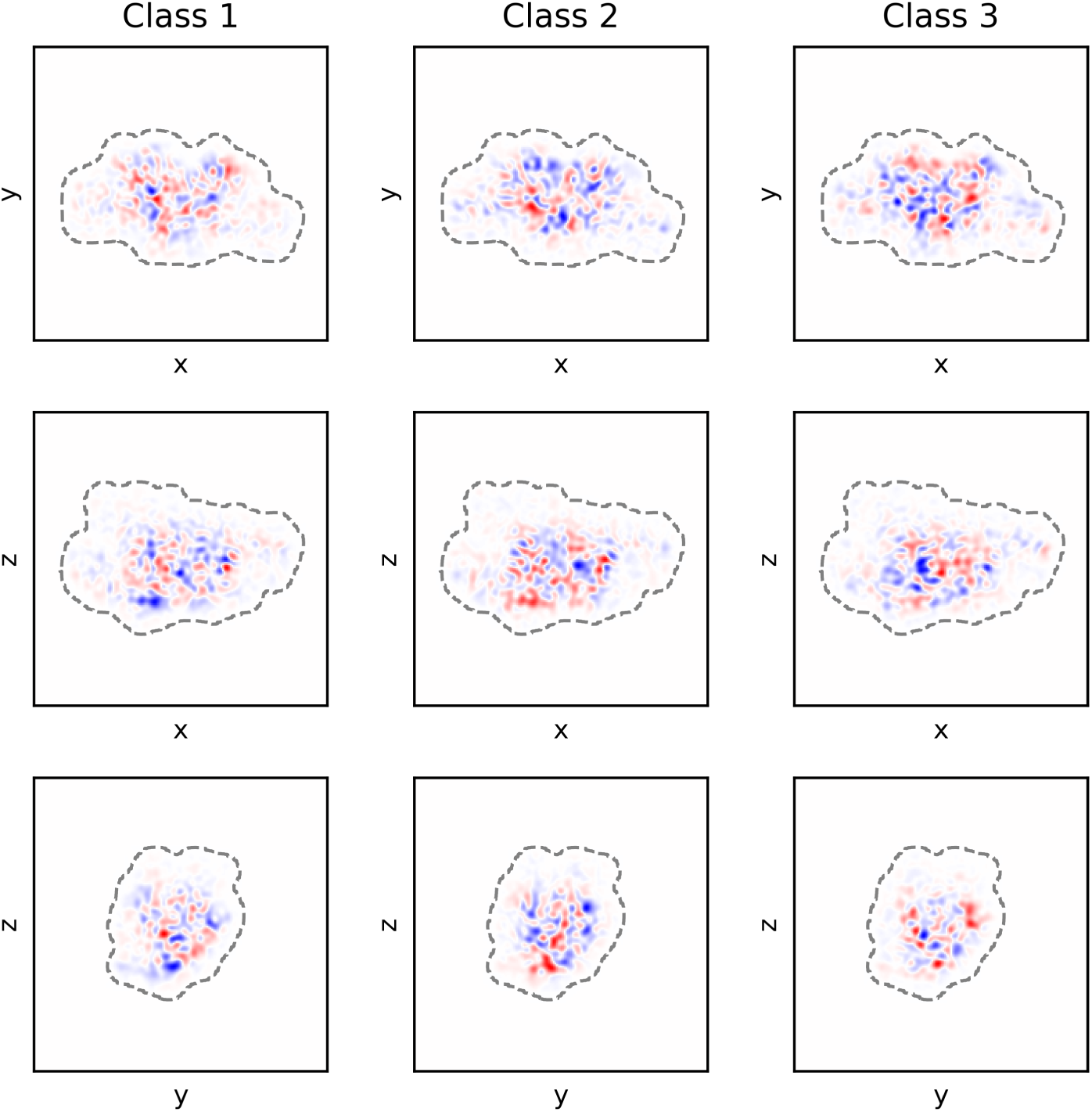
Projections of real-space difference maps of the three final density maps obtained by 3D classification. The heatmaps show projections of the difference maps by summation along each of the Cartesian axes, with blue denoting a positive sum of differences and red denoting a negative sum of differences. Grey interrupted lines illustrate the boundaries of the focus mask used during 3D classification.

**Supplementary Fig. 4:**
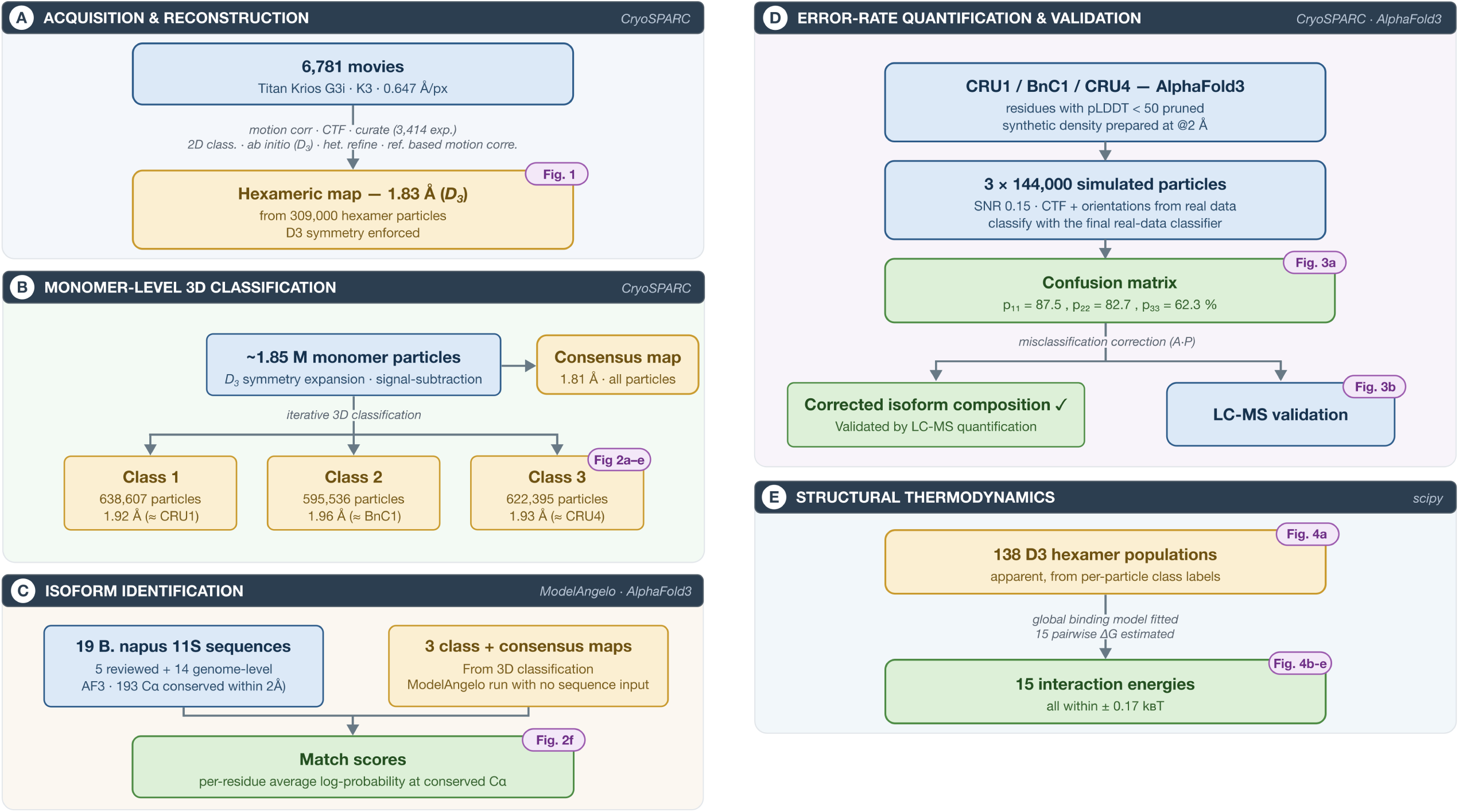
Schematic overview of the processing and analysis workflow for 3D classification of cruciferin at the monomer level and comparison to known isoform sequences. The illustrated process starts from the 309,000 particles used for the reconstruction of the homogeneous hexameric structure (Figure 1), and results in a quantitative comparison between classified monomeric particles and known cruciferin isoforms (as summarised in Figure 2f).

**Supplementary Fig. 5:**
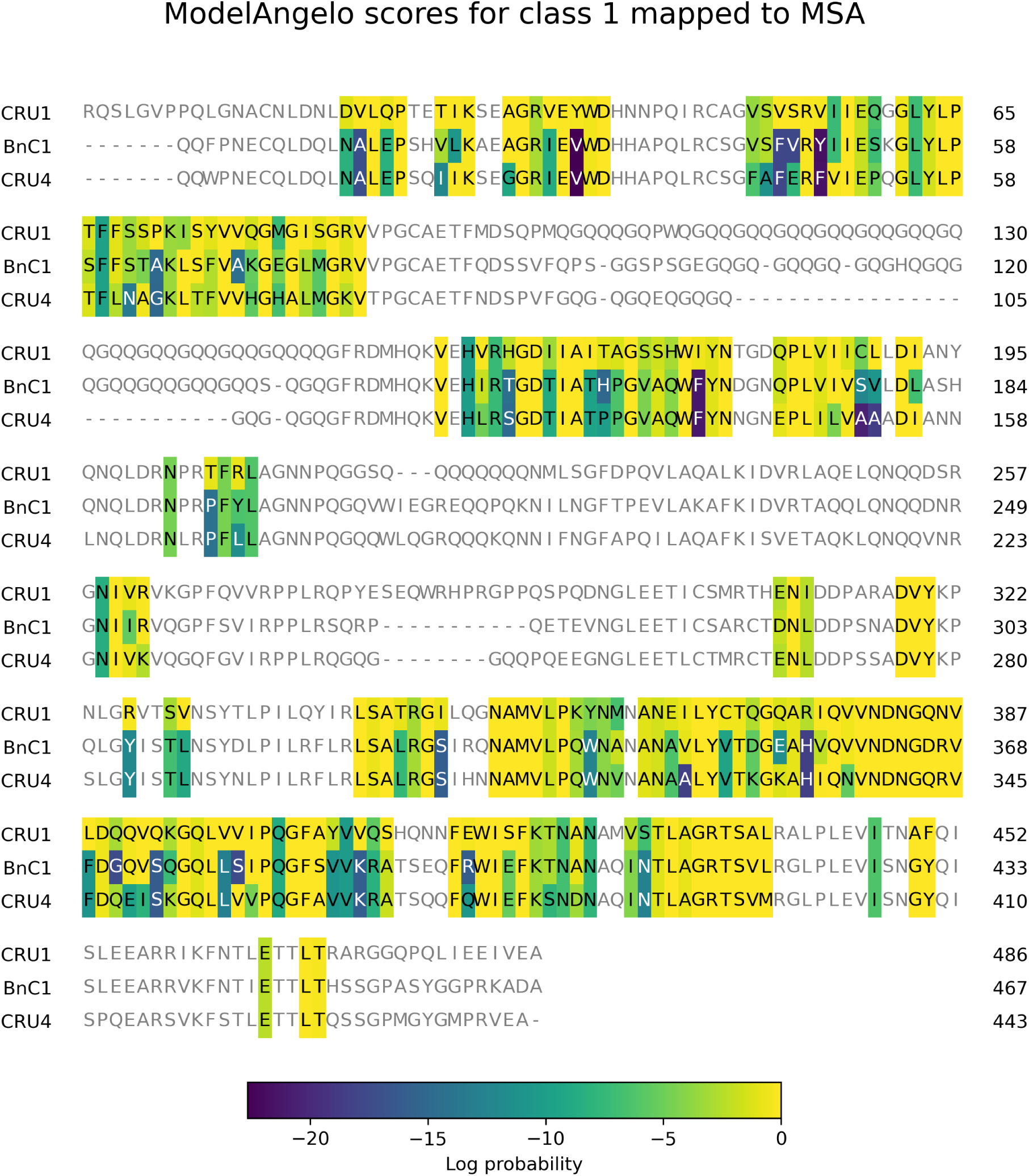
MSA of three cruciferin isoforms (CRU1, BnC1, CRU4). Residues are coloured according to the probability of a match to Class 1 from the 3D classification approach, according to ModelAngelo evalutations. Residues in grey were not matched in space between AlphaFold predictions of all 19 considered isoforms, and the chains detected by ModelAngelo.

**Supplementary Fig. 6:**
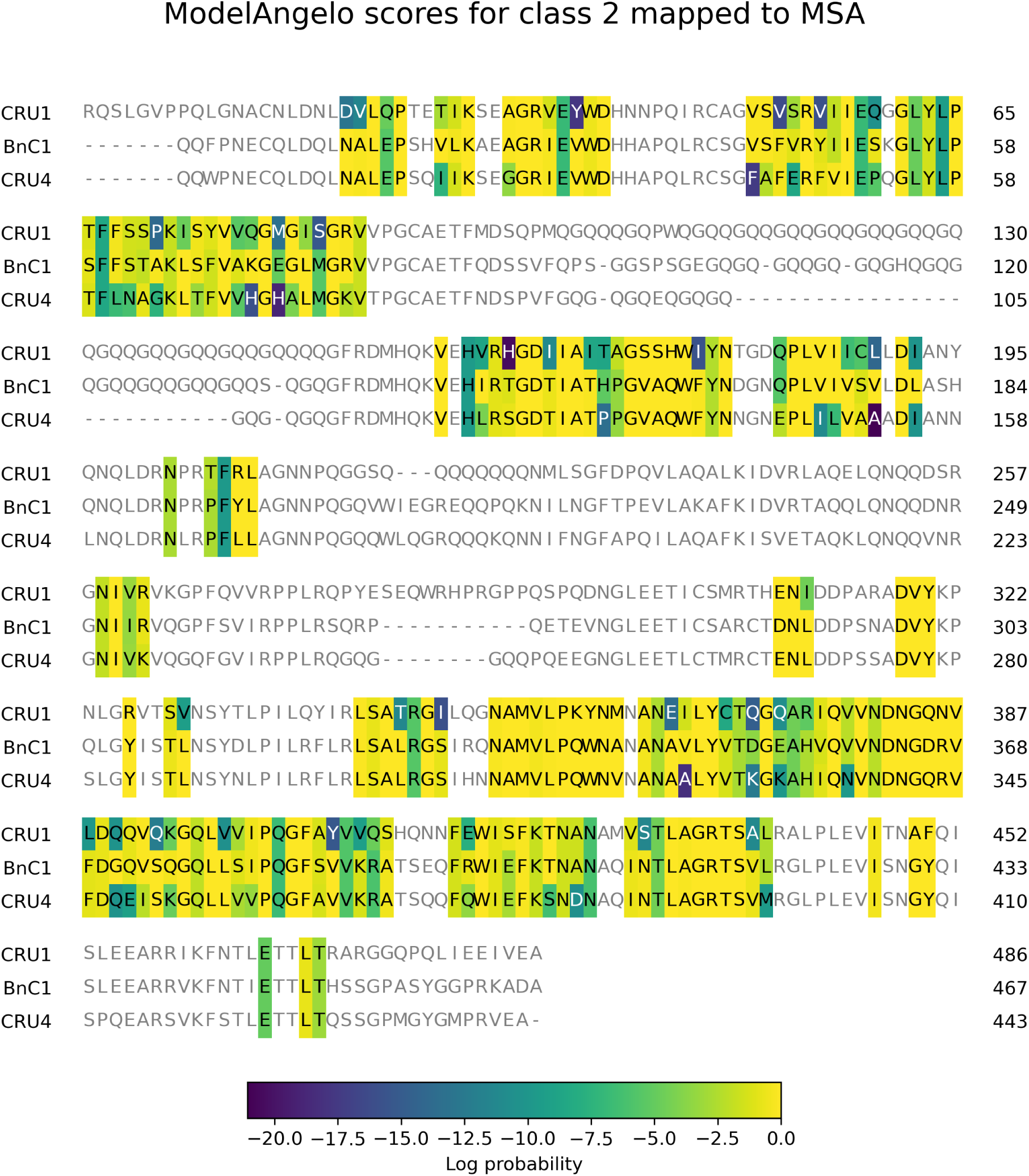
MSA of three cruciferin isoforms (CRU1, BnC1, CRU4). Residues are coloured according to the probability of a match to Class 2 from the 3D classification approach, according to ModelAngelo evalutations. Residues in grey were not matched in space between AlphaFold predictions of all 19 considered isoforms, and the chains detected by ModelAngelo.

**Supplementary Fig. 7:**
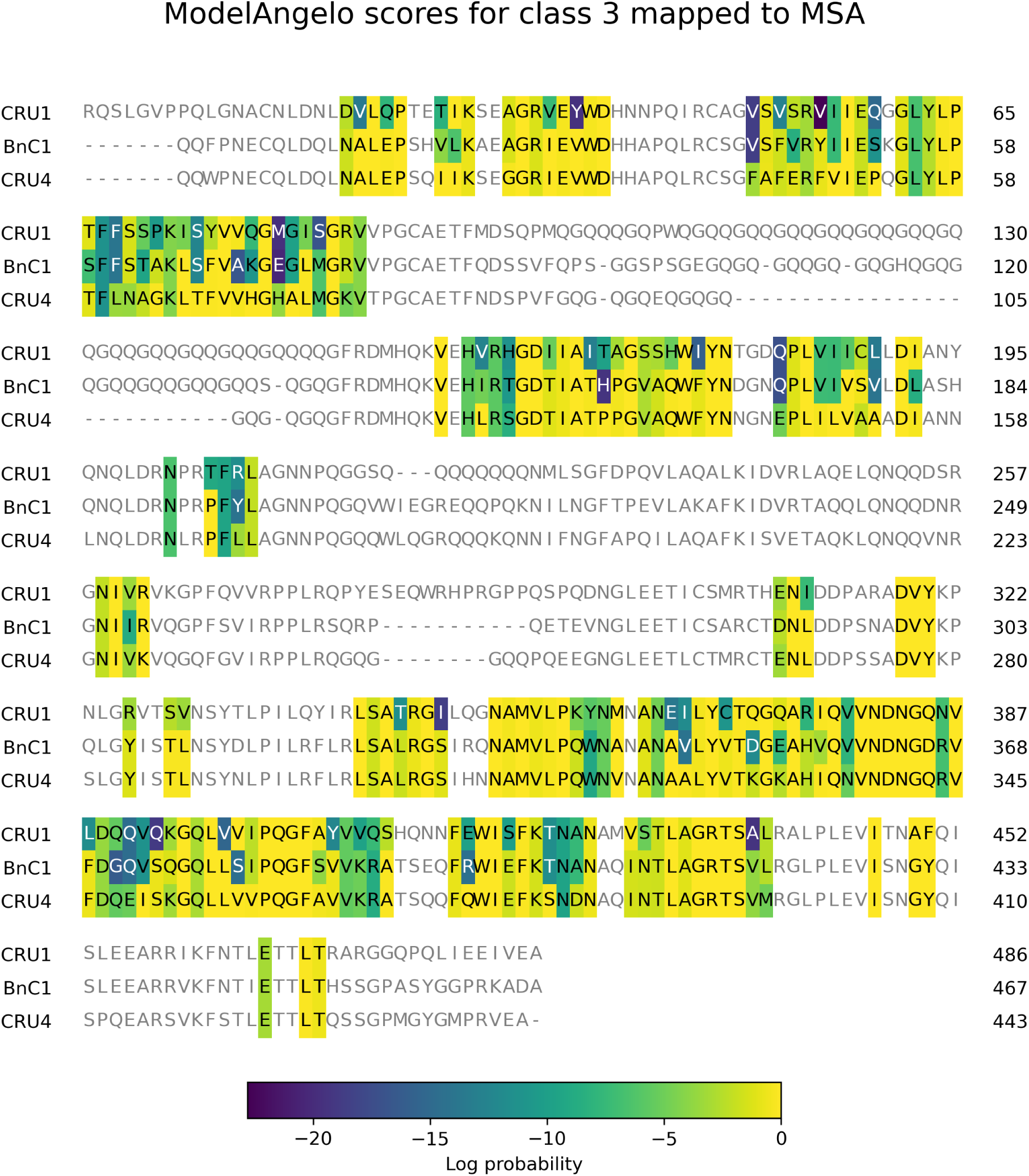
MSA of three cruciferin isoforms (CRU1, BnC1, CRU4). Residues are coloured according to the probability of a match to Class 3 from the 3D classification approach, according to ModelAngelo evalutations. Residues in grey were not matched in space between AlphaFold predictions of all 19 considered isoforms, and the chains detected by ModelAngelo.

**Supplementary Fig. 8:**
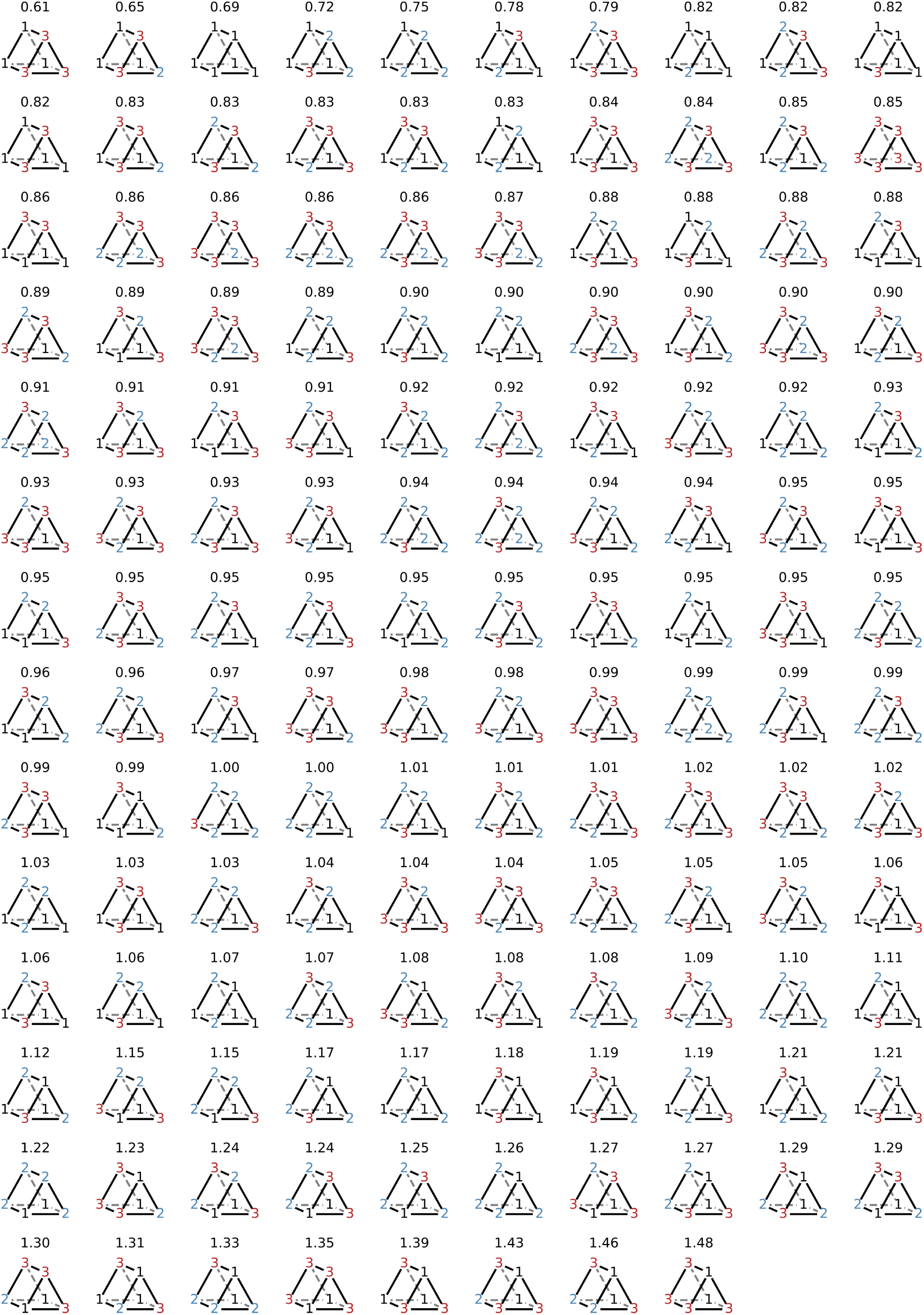
Representations of all 138 unique arrangements of three subunits in a hexamer with *D*3 symmetry, illustrated by coloured numbers at the vertices of triangular prisms. The number above each arrangement is the ratio between the apparent and expected frequencies (expected based on overall class composition and assuming no specificity, i.e., the null model).

**Supplementary Fig. 9:**
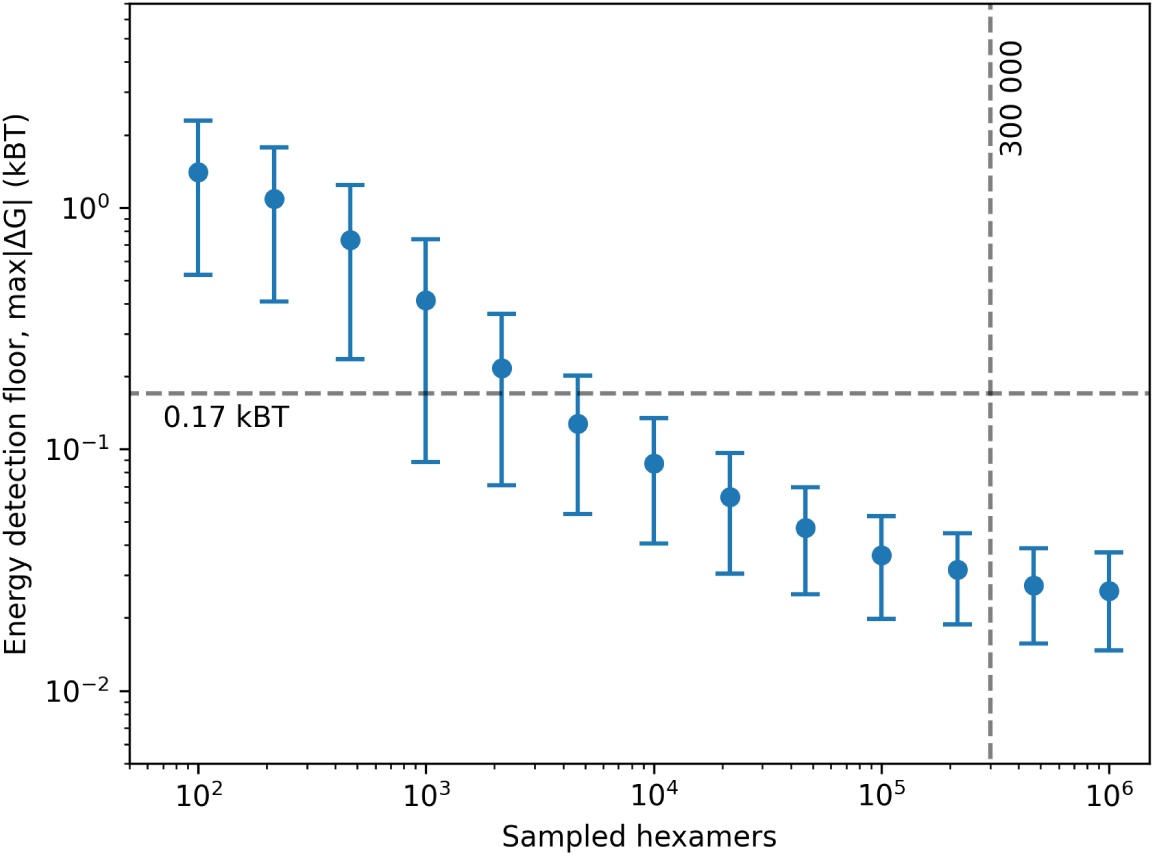
Estimated energy detection floors plotted against number of sampled hexamers. The floor is given as the average of the maximum absolute interaction energies from 1000 sampled populations based on the overall class composition and assuming no specificity in the interactions (the null model). Error bars show the standard deviation. The horizontal dashed line indicates the highest absolute value from the fit to the experimental data (0.17*k_B_ T*). The vertical dashed line indicates the 300 000 experimental hexamer particles.

**Supplementary Fig. 10:**
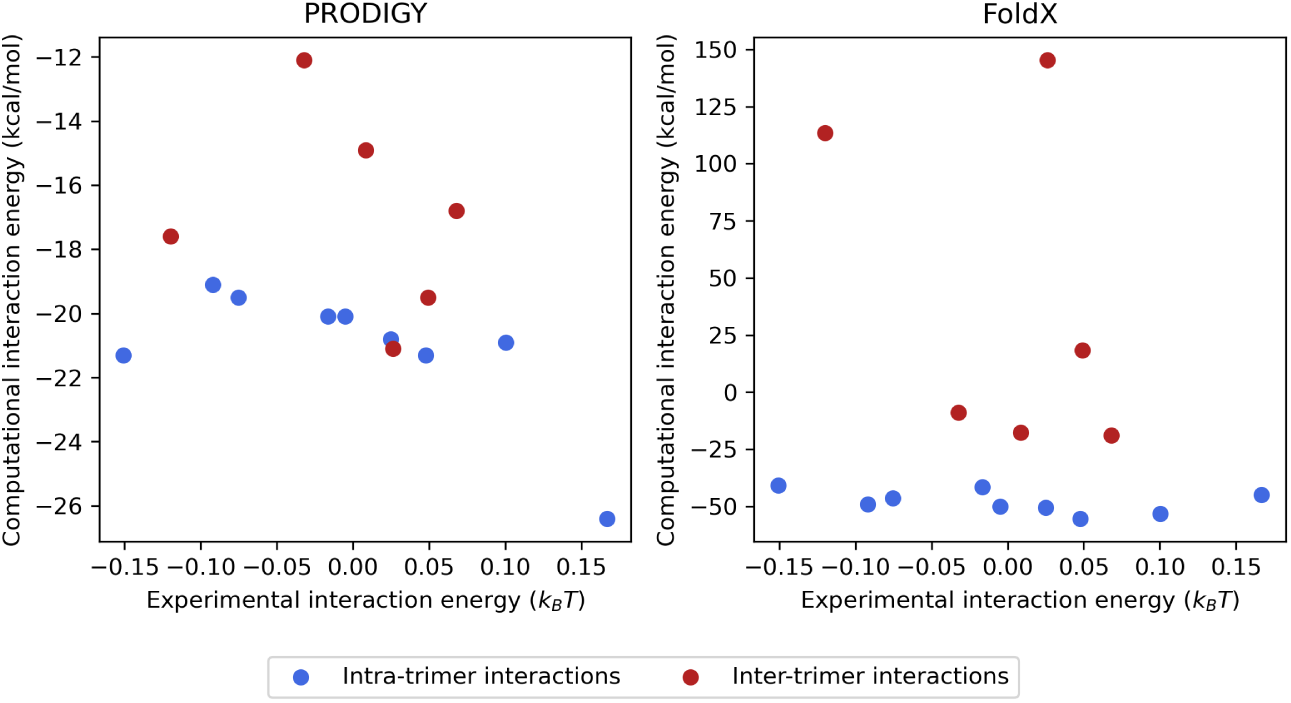
Comparison between the 15 pairwise interaction energies, as estimated both from the global binding model fitted to experimental data, and from two computational tools; PRODIGY (left) and FoldX (right).

